## Supplementary Figures 1-9 for "Pantothenate biosynthesis is critical for chronic infection by the neurotropic parasite *Toxoplasma gondii*"

### Supplementary figure 1

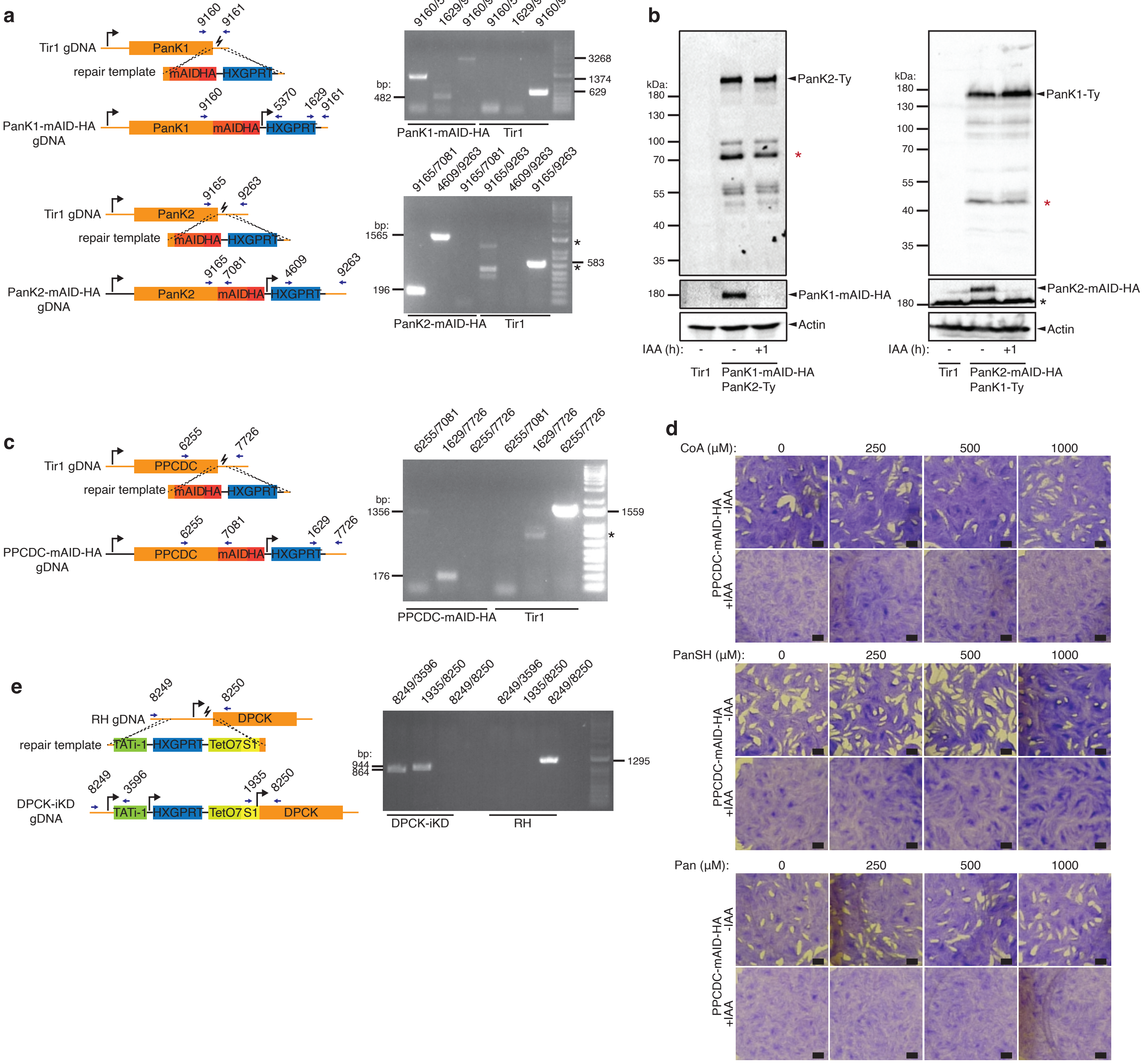

### Supplementary figure 2

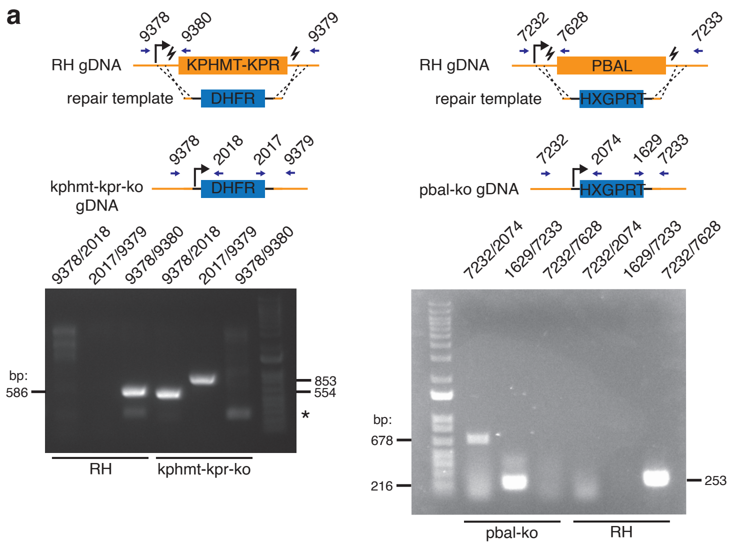


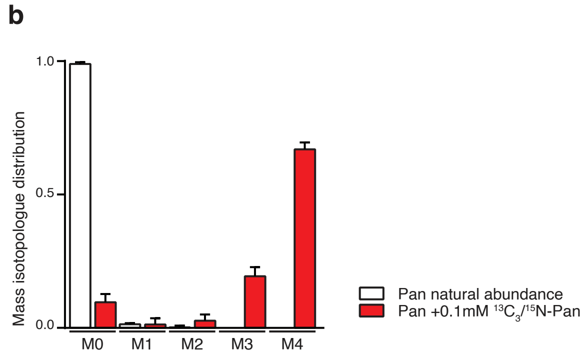

**Supplementary Fig. 3**

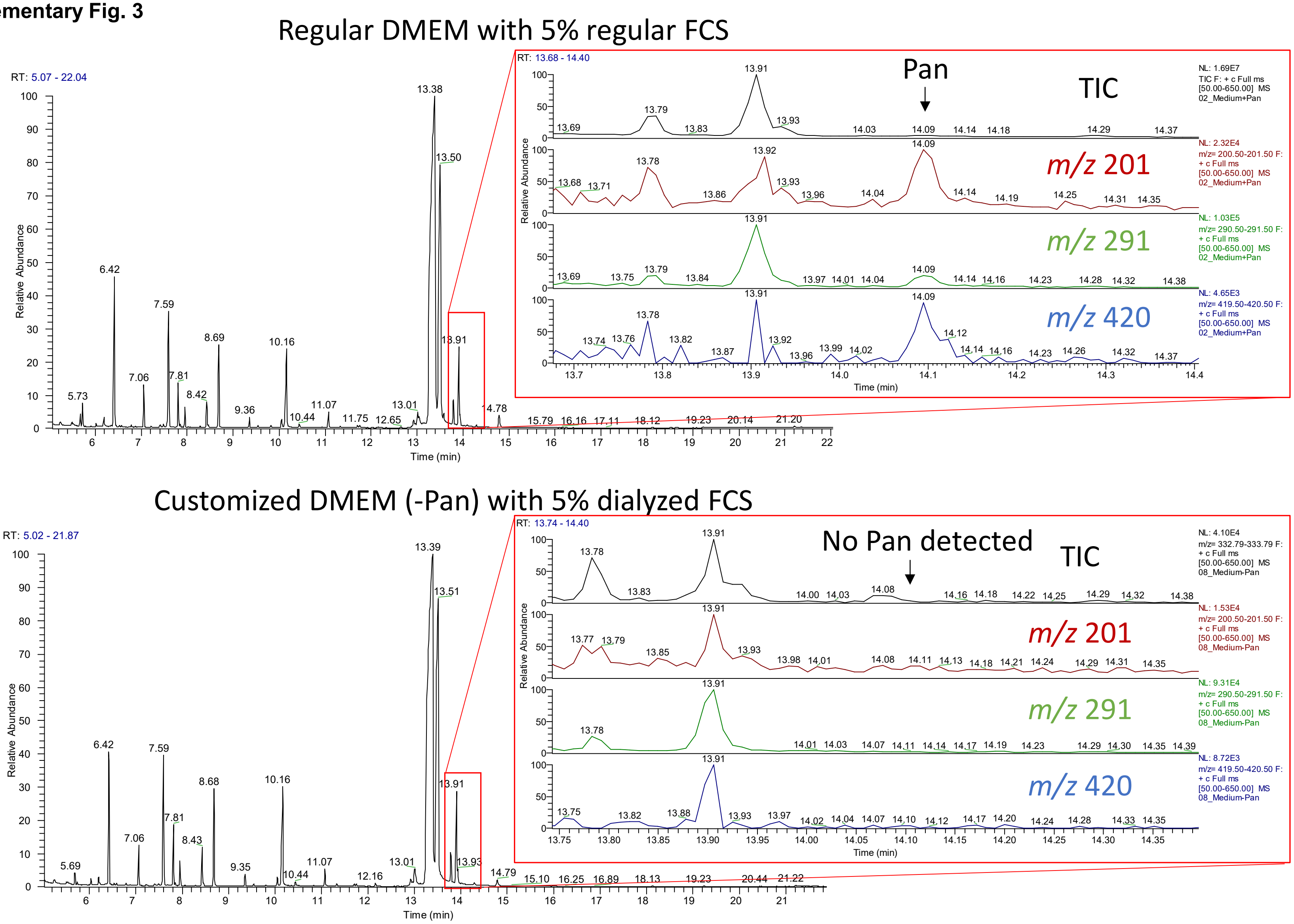

Supplementary Figure 4

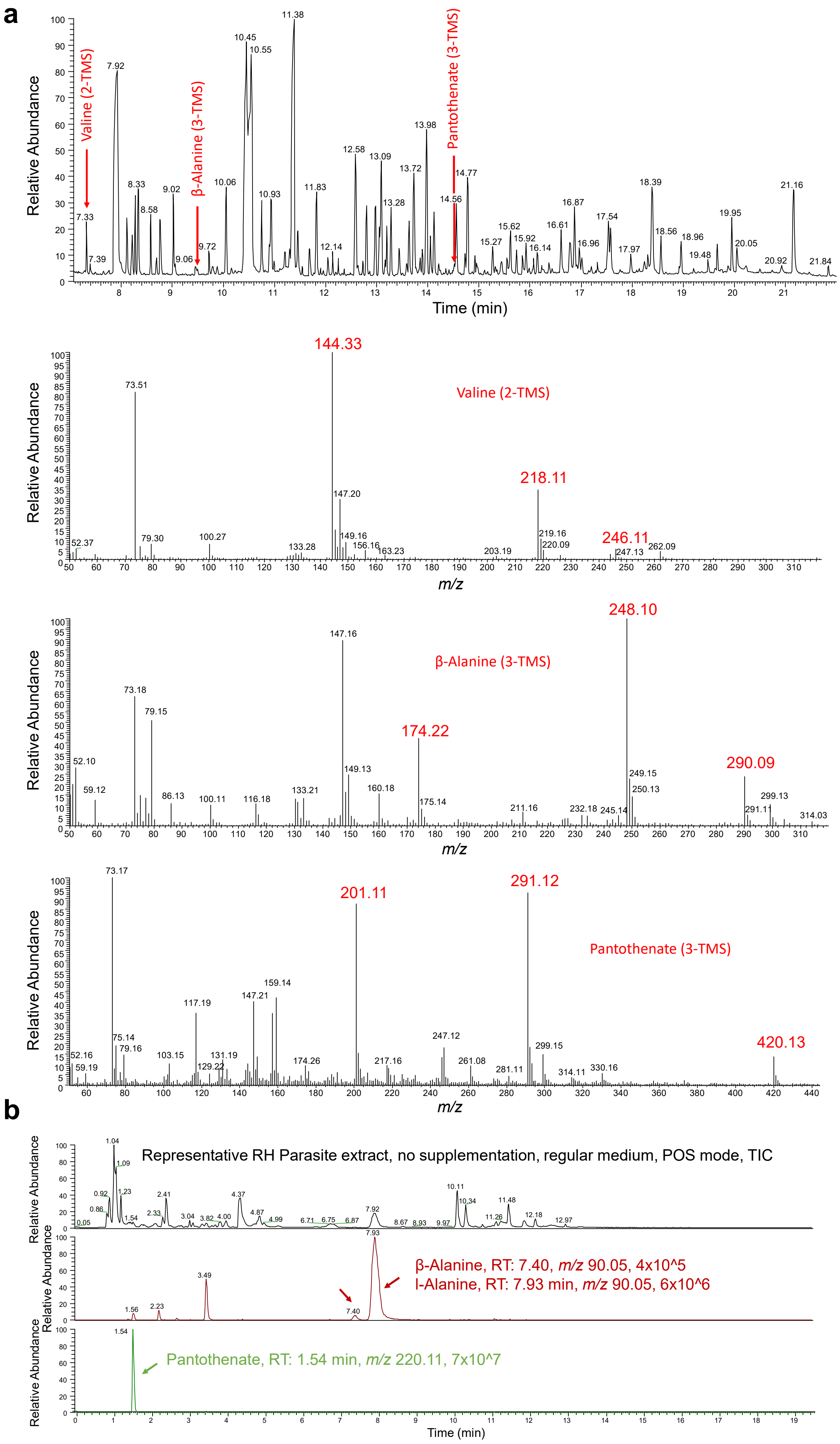

**Supplementary Figure 5**

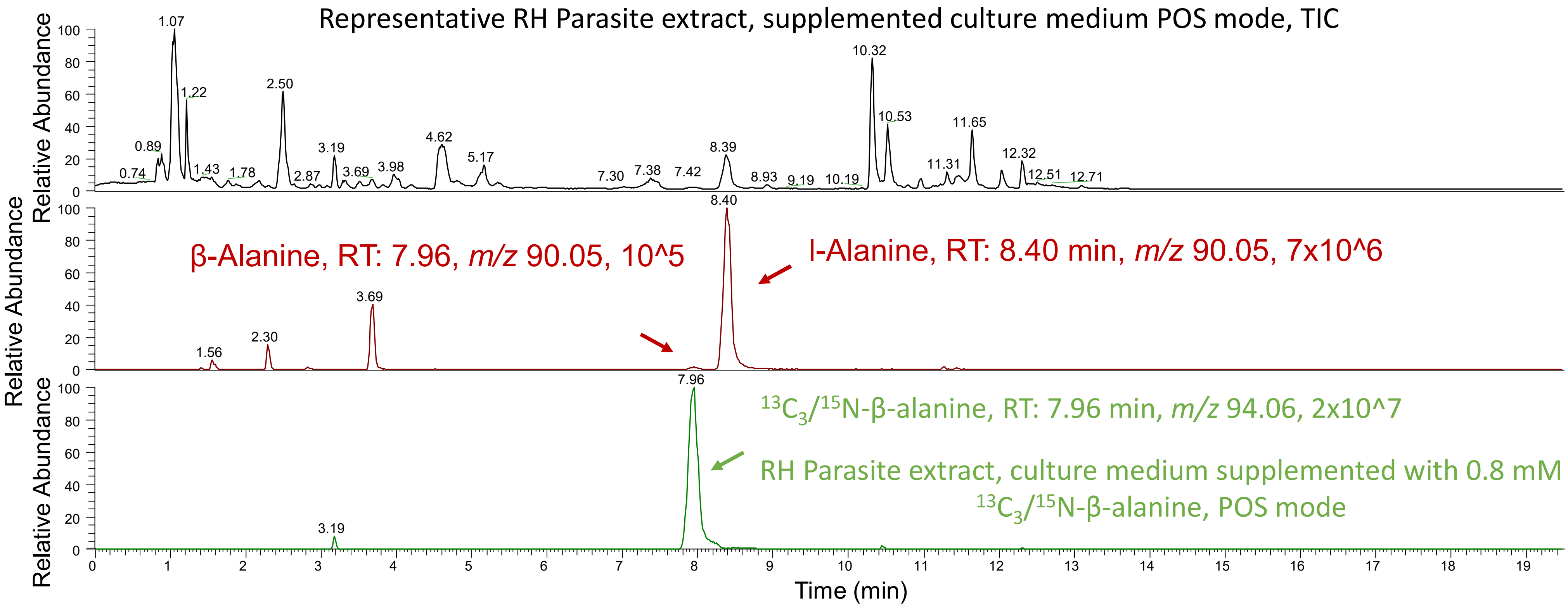

Supplementary Fig. 6

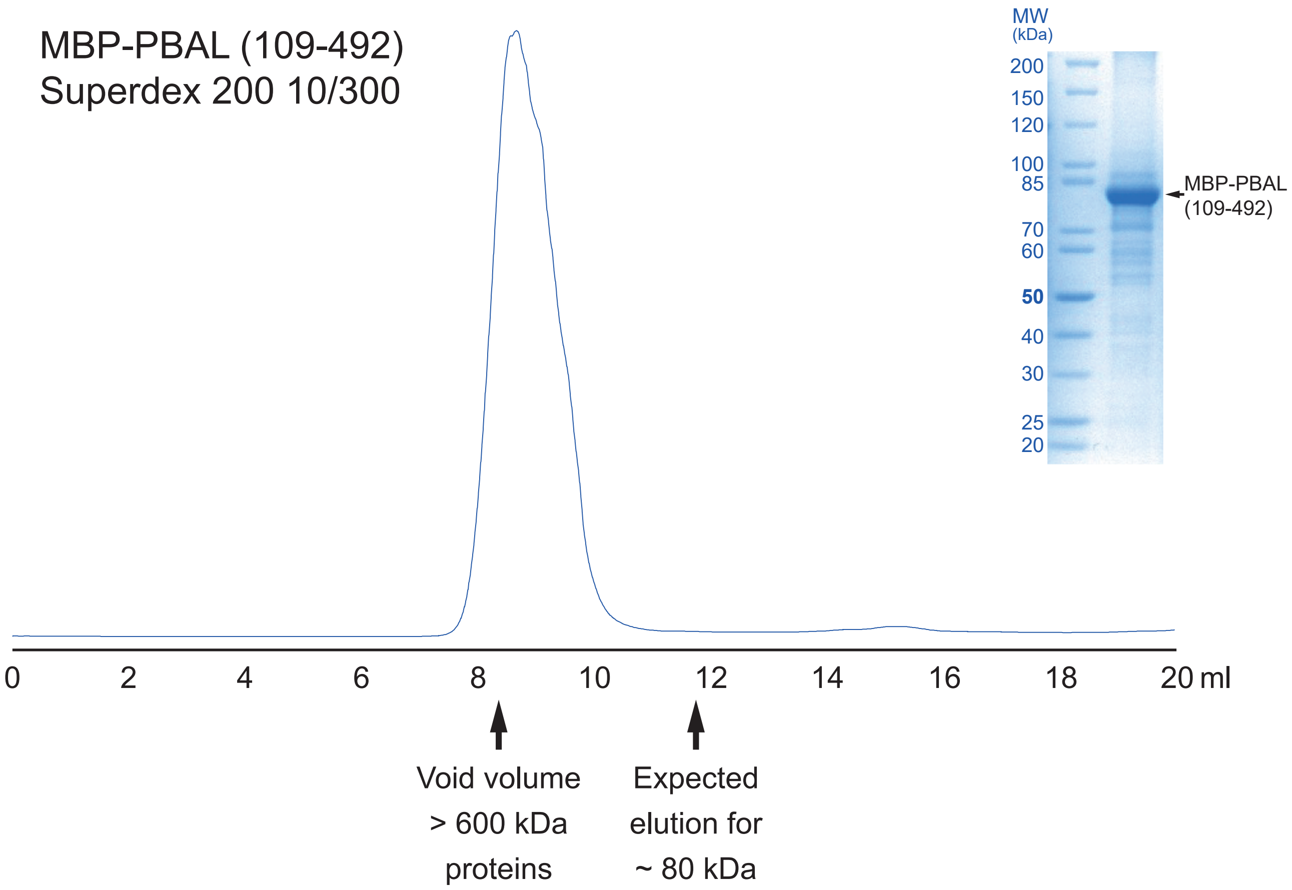

**Supplementary Figure 7**

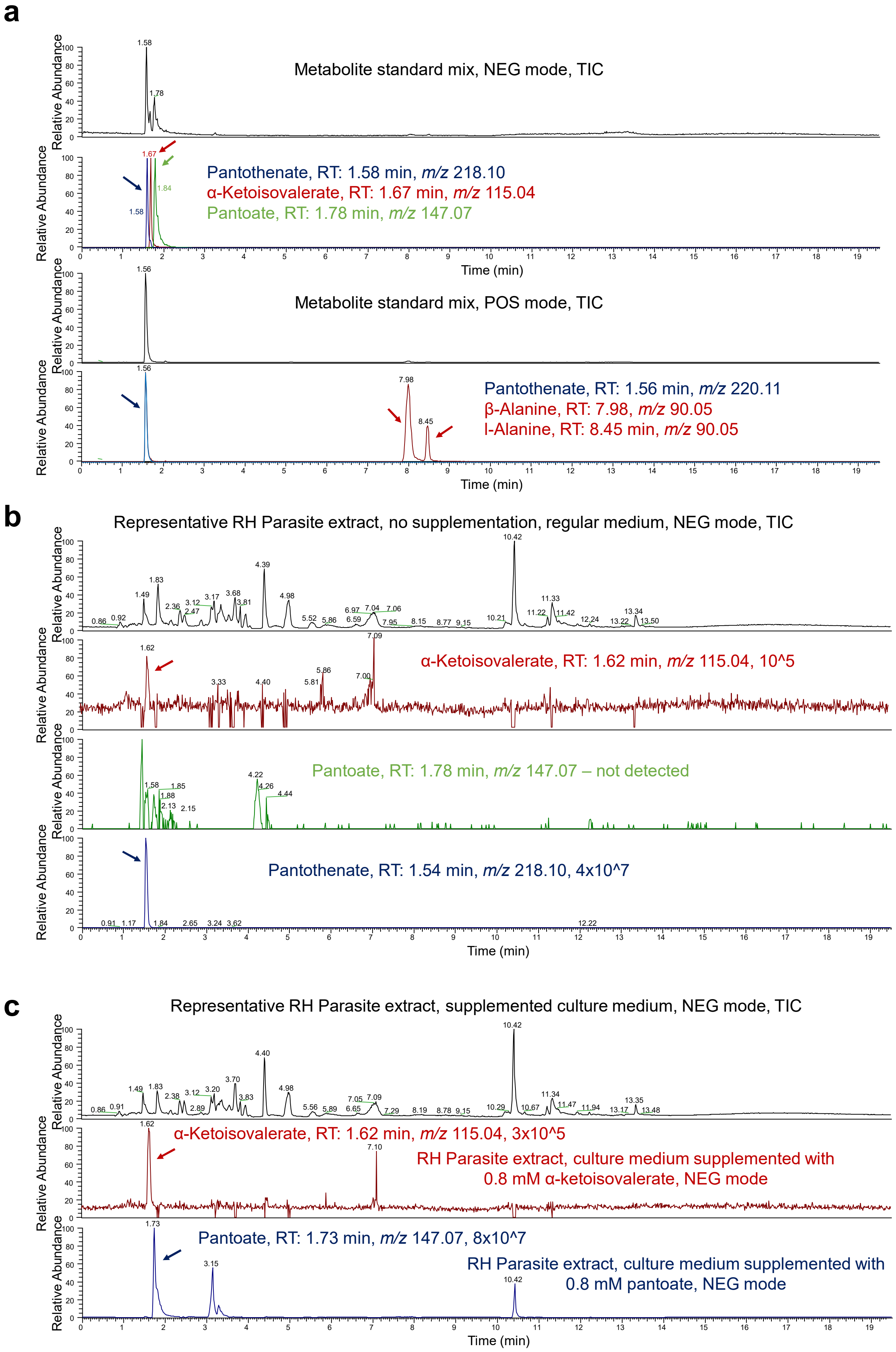

Supplementary figure 8

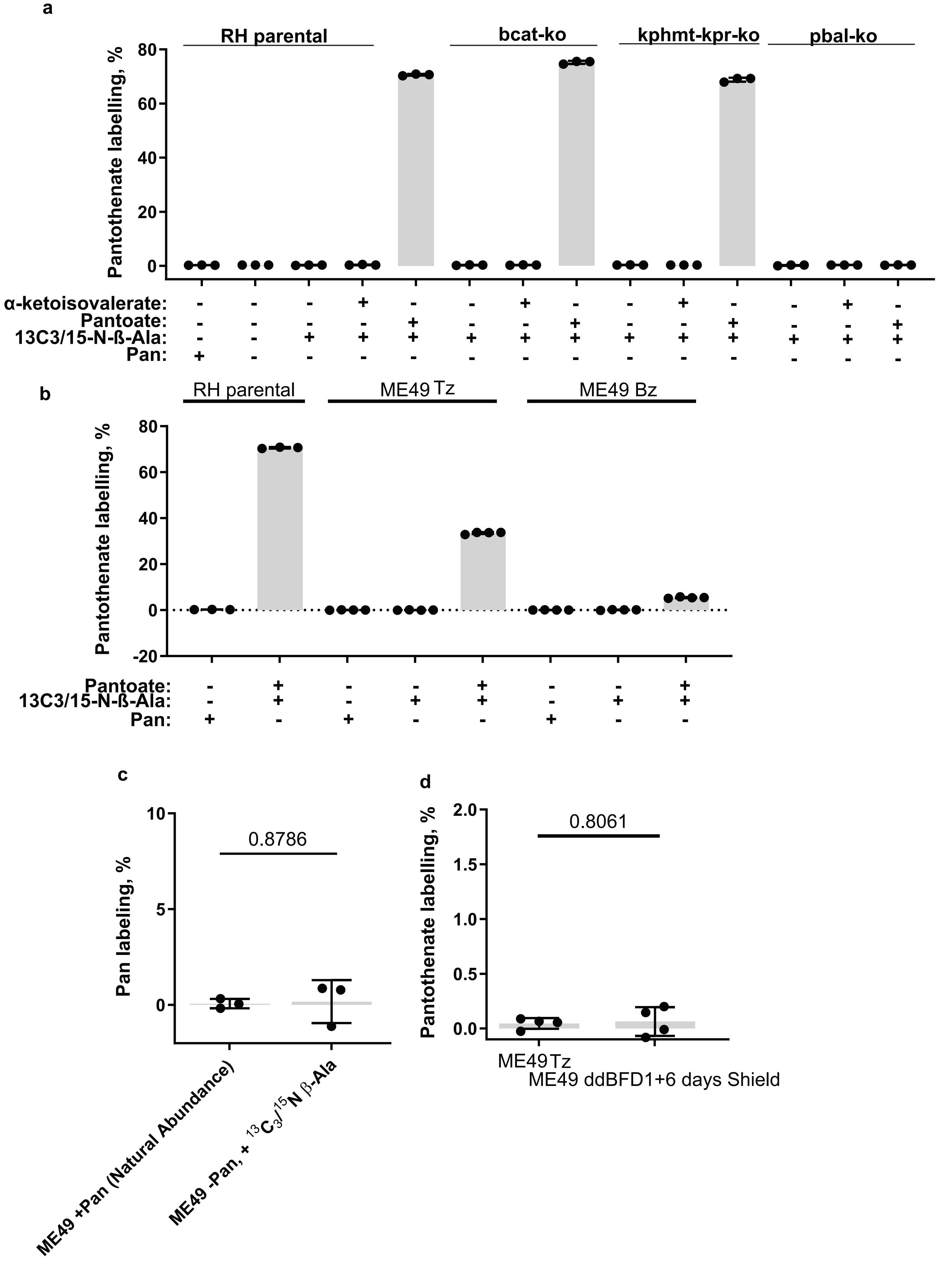

### Supplementary figure 9

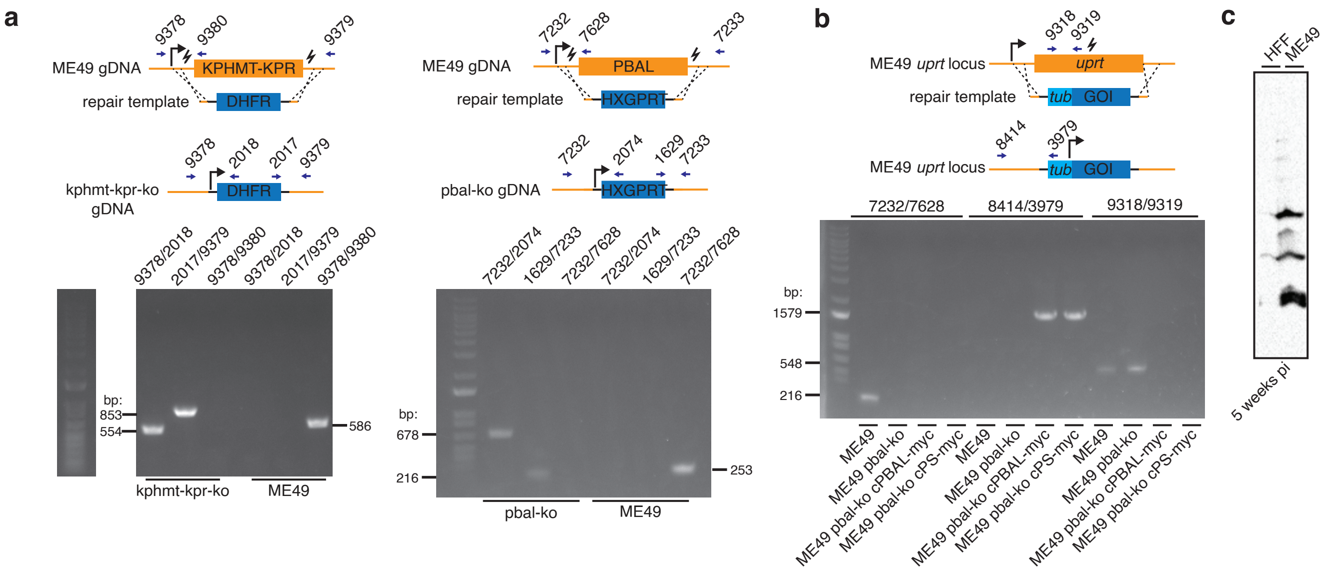
