## Supplementary File 1 for "Pantothenate biosynthesis is critical for chronic infection by the neurotropic parasite *Toxoplasma gondii*"

```
Muscle alignment of TgPBAL orthologues
Black shading represents exact match
In red catalytic residues from crystallography DOI: 10.1110/ps.0241803
In green shade if demonstrated essential for protein activity DOI: 10.1021/bi049676n


                                    10         20         30         40         50         60
                           ....|....| ....|....| ....|....| ....|....| ....|....| ....|....|
ETH_00031750               ---MNS---- ---------- ---------- ---------- ---------- ----------  3
Vbra_16581                 ---MGGVRTL YASGGGG--- ---------- -VKSLKMRDL LRKKKERRKI TVITAYDFIG  43
Cvel_3471                  MSYVGGPTSS TSEGGGASSL CFGNFMSFEP QLKSLKLEDL LGRKKAGCKI TMITAYDYLS  60
SN3_00101300               PETEETSS-- ---------- ---------- ---------- ---------- ------HQVP  12
NCLIV_039640               ---METHT-- ---------- ---------- ---------- ---------- ----------  5
TGME49_265870              ---MDTSD-- ---------- ---------- ---------- ---------- ----------  5
HHA_265870                 ---MDTSD-- ---------- ---------- ---------- ---------- ----------  5
XP_002768075.P. marinus    ---------- ---------- ---------- ---------- ---------- ----------  1
XP_002767048.P. marinus    ---------- ---------- ---------- ---------- ---------- ----------  1
sp|P31663|PANC_ECOLI       ---------- ---------- ---------- ---------- ---------- ----------  1
sp|P9WIL5|PANC_MYCTU       ---------- ---------- ---------- ---------- ---------- ----------  1

                                    70         80         90        100        110        120
                           ....|....| ....|....| ....|....| ....|....| ....|....| ....|....|
ETH_00031750               ---------- ---------- ---------- ---------- ---------P RRSEMLVLRT  14
Vbra_16581                 AQVANSAMLD CILVGDSLAN VVLGHARTNQ IGMDEMLYHC KSVTRGNSRC FVIGDMPFGS  103
Cvel_3471                  GRIADAAAID MVLVGDSLGN VVLGHSRTDK VTMEDMLHHT RAVTRGCKRA FVVGDLPFGS  120
SN3_00101300               SQALRRTETS DLLLPLGDVV LRLGSPRRVT RSSASHIHEQ AYLRRF---A SAVLPFPAVS  69
NCLIV_039640               ---------- ---------- --LAQLGRDT LP-------- ---------A FSAEFLPSSS  26
TGME49_265870              ---------- ---------- --LANVG--- -P-------- ---------Q FSVELGTRFA  22
HHA_265870                 ---------- ---------- --LANVG--- -S-------- ---------Q FSVELGTRFA  22
XP_002768075.P. marinus    ---------- ---------- ---------- ---------- ---------M LSLVPKPRSD  11
XP_002767048.P. marinus    ---------- ---------- ---------- ---------- ---------M LSLVPKPRSD  11
sp|P31663|PANC_ECOLI       ---------- ---------- ---------- ---------- ---------M LIIETLPL--  9
sp|P9WIL5|PANC_MYCTU       ---------- ---------- ---------- ---------- ---------M TIPAFHP---  8

                                   130        140        150        160        170        180
                           ....|....| ....|....| ....|....| ....|....| ....|....| ....|....|
ETH_00031750               CREVSVHRST CGRVRE---- ---------- ---------- ---------- ----------  30
Vbra_16581                 YSTVDRAVQN AIRFVNEGGV AAVKIEGPMY EQIKAVSQLT PVVGHTGVLP QTAENFLARG  163
Cvel_3471                  CCTVEKALEN AHALMRE-GC HAVKLEGPLL AQIQAISQFV PVVAHLGLQP QTASSFASRG  179
SN3_00101300               SCFPCDAVIC NTSYST---- ---------- ------MYST SATAHHGTGP WAQENWDAFG  109
NCLIV_039640               -AFF---PF- --RATA---- ---------- ---------- ---------- ----------  35
TGME49_265870              -AKFCVQRL- --NWSA---- ---------- ---------- ---------- ----------  34
HHA_265870                 -AKFCVQKL- --NLSA---- ---------- ---------- ---------- ----------  34
XP_002768075.P. marinus    ISELAAKISA RIAAKA---- ---------- ---------- ---------- ----------  27
XP_002767048.P. marinus    ISELAAKISA RITAKS---- ---------- ---------- ---------- ----------  27
sp|P31663|PANC_ECOLI       ---------- ---------- ---------- ---------- ---------- ----------  9
sp|P9WIL5|PANC_MYCTU       ---------- ---------- ---------- ---------- ---------- ----------  8

                                   190        200        210        220        230        240
                           ....|....| ....|....| ....|....| ....|....| ....|....| ....|....|
ETH_00031750               ---------- ---------- ---------- ---------- ---------- ----------  30
Vbra_16581                 KTAKDAQYLV EEAQALEAAG CCMIVLEKVC SEVAEEISRR LTIPTIGIGA GPHCDGQVLV  223
Cvel_3471                  RTADEAITLL EDARAVEAAG ASFLVLEKVA AEVAQEITSS LSIPTIGIGA GNVTDGQVLV  239
SN3_00101300               SGCLPWSRSN FPFSADCATE DAPLSTSSYM SAPSPPESGV LDACHTKAAE RPRNVTGLQK  169
NCLIV_039640               ---------- ---------- -----PPPRD R--------- ---------- ----------  41
TGME49_265870              ---------- ---------- -----NRKIC R--------- ---------- ----------  40
HHA_265870                 ---------- ---------- -----NQEIC R--------- ---------- ----------  40
XP_002768075.P. marinus    ---------- ---------- ---------- ---------- ---------- ----------  27
XP_002767048.P. marinus    ---------- ---------- ---------- ---------- ---------- ----------  27
sp|P31663|PANC_ECOLI       ---------- ---------- ---------- ---------- ---------- ----------  9
sp|P9WIL5|PANC_MYCTU       ---------- ---------- ---------- ---------- ---------- ----------  8

                                   250        260        270        280        290        300
                           ....|....| ....|....| ....|....| ....|....| ....|....| ....|....|
ETH_00031750               ---------- ---------- -------GFP ASCPILAATP EQQQQQ---- ----------  49
Vbra_16581                 WHDVLGLQSP GSLSLKFVRR FGD----GWR QTAKAVEEYA TAVERETFPH PQHNSFLMDP  279
Cvel_3471                  WHDVLGLCPP GFPKIKFAKT FGGEPGWGWK AALHAVEEYA EAVTRGIFPG -QPQTFYMGA  298
SN3_00101300               QHPCAPSQPP RTP------- -------RPI VLIHSPRELL -AYRNA---- ----------  200
NCLIV_039640               ---------- ---------- -------REI LVVHSPQELV -AYRNA---- ----------  59
TGME49_265870              ---------- ---------- -------SSH VRVLNPRNCP RNCRNG---- ----------  59
HHA_265870                 ---------- ---------- -------SSQ VRVLNPRNFP RSHRNG---- ----------  59
XP_002768075.P. marinus    ---------- ---------- -------GPP VVVRSVGDFV ALHNTD---- ----------  46
XP_002767048.P. marinus    ---------- ---------- -------GPP VVVRSVGDFV ALHNTD---- ----------  46
sp|P31663|PANC_ECOLI       ---------- ---------- ---------- --------LR QQIRRL---- ----------  17
sp|P9WIL5|PANC_MYCTU       ---------- ---------- -------GEL NVYSAPGDVA DVSRAL---- ----------  27

                                   310        320        330        340        350        360
                           ....|....| ....|....| ....|....| ....|....| ....|....| ....|....|
ETH_00031750               ---------- ---------- ---------- ---------- ---------- ----------  49
Vbra_16581                 GERDAFIRWT EGEPLSPA-- ---------- ----DVVVDQ G--------- ----------  304
Cvel_3471                  AAREDFEKRR GGRLLPPACN LRGAVGGSAS SPGVTPVSGQ GDGSTPPCSE PWRPPSTATG  358
SN3_00101300               ---------- ---------- ---------- ---------- ---------- ----------  200
NCLIV_039640               ---------- ---------- ---------- ---------- ---------- ----------  59
TGME49_265870              ---------- ---------- ---------- ---------- ---------- ----------  59
HHA_265870                 ---------- ---------- ---------- ---------- ---------- ----------  59
XP_002768075.P. marinus    ---------- ---------- ---------- ---------- ---------- ----------  46
XP_002767048.P. marinus    ---------- ---------- ---------- ---------- ---------- ----------  46
sp|P31663|PANC_ECOLI       ---------- ---------- ---------- ---------- ---------- ----------  17
sp|P9WIL5|PANC_MYCTU       ---------- ---------- ---------- ---------- ---------- ----------  27

                                   370        380        390        400        410        420
                           ....|....| ....|....| ....|....| ....|....| ....|....| ....|....|
ETH_00031750               ---------- ---------- ---------- ---------- ---------- ----------  49
Vbra_16581                 ---------- ---------- ---------- ---------- ---------- ----GEPALS  310
Cvel_3471                  RHAQAQGESA PFESVGSPSR IHGGVGLANG GGTGGVRANG GNEMLRNVAN GGSAGPPVIV  418
SN3_00101300               ---------- ---------- ---------- ---------- ---------- ----RPPVYV  206
NCLIV_039640               ---------- ---------- ---------- ---------- ---------- ----RPRV--  63
TGME49_265870              ---------- ---------- ---------- ---------- ---------- ----NTRQS-  64
HHA_265870                 ---------- ---------- ---------- ---------- ---------- ----NTRQS-  64
XP_002768075.P. marinus    ---------- ---------- ---------- ---------- ---------- ----------  46
XP_002767048.P. marinus    ---------- ---------- ---------- ---------- ---------- ----------  46
sp|P31663|PANC_ECOLI       ---------- ---------- ---------- ---------- ---------- ----------  17
sp|P9WIL5|PANC_MYCTU       ---------- ---------- ---------- ---------- ---------- ----------  27

                                   430        440        450        460        470        480
                           ....|....| ....|....| ....|....| ....|....| ....|....| ....|....|
ETH_00031750               ---------- ---------- ---------- ---------- -LLLLQQQQQ QGRCEGC---  65
Vbra_16581                 ASGVGGMVAG GLKRVLVIGS GGMASLVAYM LATKGDVAVT VATRWQQQAS AISSGGGLSC  370
Cvel_3471                  PNGPGGGMSL GFRRVAVVGC GSLGKLVGWS IAGVTGVKVS VVCRRQEQVD RILTRGGIEC  478
SN3_00101300               HHHCCGDNLH SHHNSFPSSF TRDSCDNCSD FHFDHSACPG CSPSSSKFVP PFASVGGQKY  266
NCLIV_039640               ---------- ----QLPV-- ---------- ----VAVRAA ---------- -GRTEGGAAA  82
TGME49_265870              ---------E ERQRRLPT-- ---------- ----RTAGAA LPVRARSLVN QEPNGGGPAS  99
HHA_265870                 ---------E ERQRRLPT-- ---------- ----RTVGAA LPVRARSLVI QEPNGGGPAS  99
XP_002768075.P. marinus    ---------- ---------- ---------- ---------- ---------- ----------  46
XP_002767048.P. marinus    ---------- ---------- ---------- ---------- ---------- ----------  46
sp|P31663|PANC_ECOLI       ---------- ---------- ---------- ---------- ---------- ----------  17
sp|P9WIL5|PANC_MYCTU       ---------- ---------- ---------- ---------- ---------- ----------  27

                                   490        500        510        520        530        540
                           ....|....| ....|....| ....|....| ....|....| ....|....| ....|....|
ETH_00031750               ---------- ---------- ---------- ---------- ---------- ----------  65
Vbra_16581                 VDPLGNPHG- PPVTLDVVTD DTTLPQGAFD LAIILIKSAD TERAAKLARR AVRPDGCVLS  429
Cvel_3471                  EDADGKSAGV REVTAGVASD ---FHSGSAD LVVVAVKSFD TQGAAEIAAR LCAPEGAVLS  535
SN3_00101300               VTVTDCSAS- ---------- ----TVGSVF QGDFTVGTHN VKAYQGASLE GDSCNSVLSA  311
NCLIV_039640               ADSTGASAS- ---------- ----ESGTG- ---------- ---------A GDSVDALTHA  107
TGME49_265870              ADAT----S- ---------- ----SPGRS- ---------- ---------A ATSPNASLLS  120
HHA_265870                 ADAT----S- ---------- ----SPGRS- ---------- ---------A ATWPNASLLS  120
XP_002768075.P. marinus    ---------- ---------- ---------- ---------- ---------- ----------  46
XP_002767048.P. marinus    ---------- ---------- ---------- ---------- ---------- ----------  46
sp|P31663|PANC_ECOLI       ---------- ---------- ---------- ---------- ---------- ----------  17
sp|P9WIL5|PANC_MYCTU       ---------- ---------- ---------- ---------- ---------- ----------  27

                                   550        560        570        580        590        600
                           ....|....| ....|....| ....|....| ....|....| ....|....| ....|....|
ETH_00031750               ---------- ---------- ---------- ---------- ---------- -------VAP  68
Vbra_16581                 LQNGLDAPLE LKKAF----K GRDGPKLLLG TTTNGALVRA G--------- DEGRVWRMGE  476
Cvel_3471                  LQNGLSHVGV LRKALAKSAR SAGGPCLLLG TCTAGAAPGA ADGDGSGGAV DPSRVSFRGR  595
SN3_00101300               PENEYPIEQA TEKKVSSLNS HPIRDGCPAL RVQNTSKGGL RDTCLSTRQR GYDSQSHYIR  371
NCLIV_039640               HV-------- ---------- ---------- ---------- ---------- --DRSSMFSR  117
TGME49_265870              PS-------- ---------- ---------- ---------- ---------- --SPLAFSSR  130
HHA_265870                 PS-------- ---------- ---------- ---------- ---------- --SPLASSSR  130
XP_002768075.P. marinus    ---------- ---------- ---------- ---------- ---------- -------VFR  49
XP_002767048.P. marinus    ---------- ---------- ---------- ---------- ---------- -------VFR  49
sp|P31663|PANC_ECOLI       ---------- ---------- ---------- ---------- ---------- -------RME  20
sp|P9WIL5|PANC_MYCTU       ---------- ---------- ---------- ---------- ---------- -------RLT  30

                                   610        620        630        640        650        660
                           ....|....| ....|....| ....|....| ....|....| ....|....| ....|....|
ETH_00031750               RR-------- ---------- ---------- ---------- ---------- ----------  70
Vbra_16581                 GETVIG---- --------AA ERKDEACVER VRDLLNAAAL TTRTVRSQDV LETIWHKLAI  524
Cvel_3471                  GRFVVGLMGD GEGFQRGRGG ASGPGEVAEA AASLLRSAEM DAEAVSEQEV LPALWLKVAV  655
SN3_00101300               RR-------- ---------- ---------- ---------- ---------- ----------  373
NCLIV_039640               RR-------- ---------- ---------- ---------- ---------- ----------  119
TGME49_265870              RR-------- ---------- ---------- ---------- ---------- ----------  132
HHA_265870                 RR-------- ---------- ---------- ---------- ---------- ----------  132
XP_002768075.P. marinus    GL-------- ---------- ---------- ---------- ---------- ----------  51
XP_002767048.P. marinus    GL-------- ---------- ---------- ---------- ---------- ----------  51
sp|P31663|PANC_ECOLI       GK-------- ---------- ---------- ---------- ---------- ----------  22
sp|P9WIL5|PANC_MYCTU       GR-------- ---------- ---------- ---------- ---------- ----------  32

                                   670        680        690        700        710        720
                           ....|....| ....|....| ....|....| ....|....| ....|....| ....|....|
ETH_00031750               ---------- ---------- ---------- ---------- ---------- ----------  70
Vbra_16581                 NAVVNPLTAL YGVNNGALTR PHFEPLISQI AQEVSEVGRA RGIRLPEGSN LVKVVQRAAE  584
Cvel_3471                  NAAINPLTAL MGICNGGLET QPLSTVAAAI AEEVRVVARS KGVRLSEAS- VRSALESTIR  714
SN3_00101300               ---------- ---------- ---------- ---------- ---------- ----------  373
NCLIV_039640               ---------- ---------- ---------- ---------- ---------- ----------  119
TGME49_265870              ---------- ---------- ---------- ---------- ---------- ----------  132
HHA_265870                 ---------- ---------- ---------- ---------- ---------- ----------  132
XP_002768075.P. marinus    ---------- ---------- ---------- ---------- ---------- ----------  51
XP_002767048.P. marinus    ---------- ---------- ---------- ---------- ---------- ----------  51
sp|P31663|PANC_ECOLI       ---------- ---------- ---------- ---------- ---------- ----------  22
sp|P9WIL5|PANC_MYCTU       ---------- ---------- ---------- ---------- ---------- ----------  32

                                   730        740        750        760        770        780
                           ....|....| ....|....| ....|....| ....|....| ....|....| ....|....|
ETH_00031750               ---------- ---------- ---------- ---------- ---------- ----------  70
Vbra_16581                 TTAANTSSMR TDVENRLPTE ISAICGAIAS EARKVGLSAP VNEMLVHLIK A---LEEQRQ  641
Cvel_3471                  GTARNQSSML QDVQRGSQSE VRAICGAVAE EARRNGVAAP LCSLLCLMVE GKLKLQQQPQ  774
SN3_00101300               ---------- ---------- ---------- ---------- ---------- ----------  373
NCLIV_039640               ---------- ---------- ---------- ---------- ---------- ----------  119
TGME49_265870              ---------- ---------- ---------- ---------- ---------- ----------  132
HHA_265870                 ---------- ---------- ---------- ---------- ---------- ----------  132
XP_002768075.P. marinus    ---------- ---------- ---------- ---------- ---------- ----------  51
XP_002767048.P. marinus    ---------- ---------- ---------- ---------- ---------- ----------  51
sp|P31663|PANC_ECOLI       ---------- ---------- ---------- ---------- ---------- ----------  22
sp|P9WIL5|PANC_MYCTU       ---------- ---------- ---------- ---------- ---------- ----------  32

                                   790        800        810        820        830        840
                           ....|....| ....|....| ....|....| ....|....| ....|....| ....|....|
ETH_00031750               ---------- ---------- ---------- ---------- ---------- ----------  70
Vbra_16581                 T--------- -----TPPAP ASPPLPPQPT AAKETHAALA PTAAATSPPM ATRVCKTPSE  687
Cvel_3471                  RQQQEQHSKS STYDGSSPAC VSHPSPSSNG LQANGHAEYV SAGAKVSHTA RTTLVRDPKK  834
SN3_00101300               ---------- ---------- ---------- ---------- ---------- ----------  373
NCLIV_039640               ---------- ---------- ---------- ---------- ---------- ----------  119
TGME49_265870              ---------- ---------- ---------- ---------- ---------- ----------  132
HHA_265870                 ---------- ---------- ---------- ---------- ---------- ----------  132
XP_002768075.P. marinus    ---------- ---------- ---------- ---------- ---------- ----------  51
XP_002767048.P. marinus    ---------- ---------- ---------- ---------- ---------- ----------  51
sp|P31663|PANC_ECOLI       ---------- ---------- ---------- ---------- ---------- ----------  22
sp|P9WIL5|PANC_MYCTU       ---------- ---------- ---------- ---------- ---------- ----------  32

                                   850        860        870        880        890        900
                           ....|....| ....|....| ....|....| ....|....| ....|....| ....|....|
ETH_00031750               ---------- --RIGFVPTM GGLHWGHLSL IRAAERH--- CDEVWVSIFL NPTQFQSQED  115
Vbra_16581                 LMGVRSTLPA SARVAFVPFL GGLHDGHMAL LDEASRR--- GDRVWTSLFL NQLQFQSAKD  744
Cvel_3471                  LTALRKQVPP SSRVVFVPFL GGLHHGHLAL VEDARRRCGE GGQVWCSVFL NRLQFQSAQD  894
SN3_00101300               ---------- --RISLVPTL GGLHEGHLHL MRQAACL--- SDEVWVSIFL NPLQFQSSKD  418
NCLIV_039640               ---------- --RISLVPTM GGLHEGHLHL IRGAACT--- GDEVWVTIFV NALQFHSAKD  164
TGME49_265870              ---------- --RISLVPTM GGLHEGHLHL IRGAACT--- GDEVWVTIFV NALQFHSAKD  177
HHA_265870                 ---------- --RISLVPTM GGLHEGHLHL IRGAACT--- GDEVWVTIFV NALQFHSAKD  177
XP_002768075.P. marinus    ---------- --SVGFVPTM GSLHSGHMKL IATARPN--- HDVLVVSIFV NPAQFSPEED  96
XP_002767048.P. marinus    ---------- --SVGFVPTM GSLHSGHMKL IATARPN--- HDVLVVSIFV NPAQFSPEED  96
sp|P31663|PANC_ECOLI       ---------- --RVALVPTM GNLHDGHMKL VDEAKAR--- ADVVVVSIFV NPMQFDRPED  67
sp|P9WIL5|PANC_MYCTU       ---------- --RVMLVPTM GALHEGHLAL VRAAKRV--P GSVVVVSIFV NPMQFGAGED  78

                                   910        920        930        940        950        960
                           ....|....| ....|....| ....|....| ....|....| ....|....| ....|....|
ETH_00031750               FDTYPIKVEE DLHLLTTQ-- TNAQVLFLPE ASEMYPHLAP RRGAPAGAPG GGPPVGAP--  171
Vbra_16581                 FETYPMRLDE DIAKLRDR-- N-VEVIFTPN VQDIYP--TA EDSF------ ----------  783
Cvel_3471                  FDTYPMDLQR DVQMLEEA-- G-VDVVFAPS SEGVYS--DK DAVF------ ----------  933
SN3_00101300               LLTYPASIDE DIRLIQQL-- GVVSVLFVPH REALFP-LDA PAGF---SAS PGVTSASASS  472
NCLIV_039640               FLTYPASIDD DMKLLSQV-- R-VDLVFIPS HSSLYP-LDS RPHL---IAG SEELYTVP--  215
TGME49_265870              FLTYPASIDD DMKLLSQV-- R-VDLVFIPS HSSLYP-LDR RPHL---IAG SEVLSTIPAR  230
HHA_265870                 FLTYPASIDD DMKLLSQV-- R-VDLVFIPS HSSLYP-LDR RPHL---IAG SEVLSTIPAR  230
XP_002768075.P. marinus    YEQYPRDLEG DLKKLETESA G-VDVVFAPE PADMYP-KNP RAIV------ ----------  138
XP_002767048.P. marinus    YEQYPRDLEG DLKKLETESA G-VDVVFAPE PADMYP-KNP RAIV------ ----------  138
sp|P31663|PANC_ECOLI       LARYPRTLQE DCEKLNKR-- K-VDLVFAPS VKEIYP-NGT ETHT------ ----------  107
sp|P9WIL5|PANC_MYCTU       LDAYPRTPDD DLAQLRAE-- G-VEIAFTPT TAAMYP-DGL RTTV------ ----------  118

                                   970        980        990        1000       1010       1020
                           ....|....| ....|....| ....|....| ....|....| ....|....| ....|....|
ETH_00031750               GEGLAPGGRP TAGGPPVGAP AQRRGVPFAE ---------- ---------- ----------  201
Vbra_16581                 ---------- ---------- ---------- ---------- ---------- ----------  783
Cvel_3471                  ---------- ---------- ---------- ---------- ---------- ----------  933
SN3_00101300               EHGIRAGHLT ACGERKESTS EEGRETASQD SETLVSVPPA AAACHKTAAA VGDSAGTVTP  532
NCLIV_039640               -RGRDNG--- --DARANGLP NATD----KE ADRVVRSPVS GFKLERLDTH AGERDA----  261
TGME49_265870              GRGLEDG--- --GGRVTSGQ SDDEADAMEE ADRVELSPVD --RLNRGDAR AREVDG----  279
HHA_265870                 GRGLEDG--- --GGRVTSGQ SDDEADAMEE ADRVEVSPVD --RLNRGDAR GREVDG----  279
XP_002768075.P. marinus    ---------- ---------- ---------- ---------- ---------- ----------  138
XP_002767048.P. marinus    ---------- ---------- ---------- ---------- ---------- ----------  138
sp|P31663|PANC_ECOLI       ---------- ---------- ---------- ---------- ---------- ----------  107
sp|P9WIL5|PANC_MYCTU       ---------- ---------- ---------- ---------- ---------- ----------  118

                                   1030       1040       1050       1060       1070       1080
                           ....|....| ....|....| ....|....| ....|....| ....|....| ....|....|
ETH_00031750               ---------- ---------- ---------- ---------- ---------- ----------  201
Vbra_16581                 ---------- ---------- ---------- ---------- ---------- ----------  783
Cvel_3471                  ---------- ---------- ---------- ---------- ---------- ----------  933
SN3_00101300               SVKLEGSSEL FLQGNDGDRL PSKVAATLGS VVYDKASPAR SVLTATSTPT AEESSAHEKG  592
NCLIV_039640               ---------- ---------- ---------- ---LGPTPA- ----ASWIP- ----------  272
TGME49_265870              ---------- ---------- ---------- ---DGCSHS- ----A-WSP- ----------  289
HHA_265870                 ---------- ---------- ---------- ---DDCSHS- ----A-WSP- ----------  289
XP_002768075.P. marinus    ---------- ---------- ---------- ---------- ---------- ----------  138
XP_002767048.P. marinus    ---------- ---------- ---------- ---------- ---------- ----------  138
sp|P31663|PANC_ECOLI       ---------- ---------- ---------- ---------- ---------- ----------  107
sp|P9WIL5|PANC_MYCTU       ---------- ---------- ---------- ---------- ---------- ----------  118

                                   1090       1100       1110       1120       1130       1140
                           ....|....| ....|....| ....|....| ....|....| ....|....| ....|....|
ETH_00031750               ---------- ---------- -LVVDFDGIE EVEGEGKRRP GFFR------ ----------  224
Vbra_16581                 ---------- ---------- PARVDFAGIE AVDGEGKERP GFFRGIGTVL TKFFAWTRPS  823
Cvel_3471                  ---------- ---------- PARIDFEGVE SVGGEGNMRP GFFKGVGTVV GKFFAWVRPD  973
SN3_00101300               NSGRRRQQEK PNSSGSSRLF RMRVFFKDIE DVQGEGRRRR GFFGGIGTVV AQLLTLTRPT  652
NCLIV_039640               ---------- -----PQRLF RMRVDFEGIE EVEGEGRRRS GFLRGIGTVV TQLFTLIRPT  317
TGME49_265870              ---------- -----PQRLF RMRVDFEGIE DVEGEGRRRC GFFRGIGTVV VQLFTLIRPT  334
HHA_265870                 ---------- -----PQRLF RMRVDFEGIE DVEGEGRRRC GFFRGIGTVV VQLFTLIRPT  334
XP_002768075.P. marinus    ---------- ---------- PSVTVEPNFV NGLSEAACRP TFFRGVATVV MKLFNIIRPE  178
XP_002767048.P. marinus    ---------- ---------- PSVTVEPNFV NGLSEAACRP TFFRGVATVV MKLFNIIRPE  178
sp|P31663|PANC_ECOLI       ---------- ---------- --YVDVPGLS TML-EGASRP GHFRGVSTIV SKLFNLVQPD  144
sp|P9WIL5|PANC_MYCTU       ---------- ---------- -----QPGPL AAELEGGPRP THFAGVLTVV LKLLQIVRPD  153

                                   1150       1160       1170       1180       1190       1200
                           ....|....| ....|....| ....|....| ....|....| ....|....| ....|....|
ETH_00031750               ---------- --------VS SGCIDTRVVL CPTARG-PQG LPFASRNKRL TQQQLEKAQE  265
Vbra_16581                 VVVFGQKDFL QTVIVRKLCA EFFPDVEVAV VPTVRE-KDG LAFASRNARL TPEERKRAPL  882
Cvel_3471                  VAIFGQKDFL QTVVVKRLAR EFFIDTEIVV HETVRE-ESG LATASRNEKL TKEERHKAAG  1032
SN3_00101300               YVHVGLKDFQ QVICLKRLVC GLGLDVLVVA HSTKRHASDG LSLSSRNKRL SGAERTAAAK  712
NCLIV_039640               YAHFGFKDFQ QVACVKRLIC GMGLDALLLA HNTWRD-SDG MAASSRNRRM TEEDRKKARK  376
TGME49_265870              YAHCGFKDFQ QVACLKRLIC GMGLDALLLA HNTWRD-SDG VAASSRNRRM AEADRQKARK  393
HHA_265870                 YAHCGFKDFQ QVACLKRLIC GMGLDALLLA HNTWRD-SDG VAASSRNRRM AEADRQKARK  393
XP_002768075.P. marinus    RAYFGQKDAM QVSVIISMVK DLNVPVELEV VPTARE-ADG LASSSRNVYL TPAMREKATI  237
XP_002767048.P. marinus    RAYFGQKDAM QVSVIISMVK DLNVPVELEV VPTARE-ADG LASSSRNVYL TPAMREKATI  237
sp|P31663|PANC_ECOLI       IACFGEKDFQ QLALIRKMVA DMGFDIEIVG VPIMRA-KDG LALSSRNGYL TAEQRKIAPG  203
sp|P9WIL5|PANC_MYCTU       RVFFGEKDYQ QLVLIRQLVA DFNLDVAVVG VPTVRE-ADG LAMSSRNRYL DPAQRAAAVA  212

                                   1210       1220       1230       1240       1250       1260
                           ....|....| ....|....| ....|....| ....|....| ....|....| ....|....|
ETH_00031750               ACAVLKEVEA LYNKGTR-QI GALRAAA--- AAAAERVG-A HLEYLAFNRQ ADGAPIVPLG  320
Vbra_16581                 IYQTLQAVEA QHRRGER-DA GVLTEAG--- RVFAASHG-L SVDYITVCDL YTGKAADVLQ  937
Cvel_3471                  LFASLSLVKD AVESGEVTSV SDLISLG--- LTALKKRE-M HMDYLILSDL ESGRELQEDE  1088
SN3_00101300               YYSVLQQVCR SYERGER-RV QQLETEA--- RLHAAAAG-L HLYYITFNRW QDGAPVAYTS  767
NCLIV_039640               SFLLLQHVSR -----ER--- ---------- ---------- ---------- ----------  388
TGME49_265870              SFLLLQHVGE VFRKGER-RV HRLLEEA--- RAFAEKEN-L LLHYTAIDRL EDGEPVVYRH  448
HHA_265870                 SFLLLQHVGE VFRKGER-RV HTLLEEA--- RAFAEKEN-L LLHYTAIDRL EDGEPVVYRH  448
XP_002768075.P. marinus    LYKSLCAAYD MVKSASK-PV KASEVEDVVK RTLLAEPMVL GIEYISVASV ETAQEVETIQ  296
XP_002767048.P. marinus    LYKSLCAAYD MVKSASK-PV KASEVEDVVK RTLLAEPMVL GIEYISVASV ETAQEVETIQ  296
sp|P31663|PANC_ECOLI       LYKVLSSIAD KLQAGER-DL DEIITIA--- GQELNEKG-F RADDIQIRDA DTLLEVSETS  258
sp|P9WIL5|PANC_MYCTU       LSAALTAAAH AATAGAQ--- -AALDAA--- RAVLDAAPGV AVDYLELRDI GLG-PMPLNG  264

                                   1270       1280       1290       1300       1310       1320
                           ....|....| ....|....| ....|....| ....|....| ....|....| ....|....|
ETH_00031750               APRGAPRR-- ---------- ---------- ---------- ---------- -ASEGAPGGT  337
Vbra_16581                 PT-------- ---------- ---------- ---------- ---------- ----------  939
Cvel_3471                  EVCPGAAL-- ---------- ---------- ---------- ---------- ----------  1096
SN3_00101300               GSSWTRKTPS CCSATSNTTD SVAPASCSSL GANQQEHTRQ QEQPAQQQLA RRKEDAEAQT  827
NCLIV_039640               ---------- ---------- ---------- ---------- ---------- ----------  388
TGME49_265870              SGGAVKRIET QGESAEN--- ---------- --EERRAKRR RTESDIADET PLGRPGTGSP  493
HHA_265870                 SGGAVKRIDT HRESAAN--- ---------- --GERRAKRR RTDSDIADEA QLGRPGTGSP  493
XP_002768075.P. marinus    FGPEA----- ---------- ---------- ---------- ---------- ----------  301
XP_002767048.P. marinus    FGPEA----- ---------- ---------- ---------- ---------- ----------  301
sp|P31663|PANC_ECOLI       ---------- ---------- ---------- ---------- ---------- ----------  258
sp|P9WIL5|PANC_MYCTU       SGR------- ---------- ---------- ---------- ---------- ----------  267

                                   1330       1340       1350       1360       1370       1380
                           ....|....| ....|....| ....|....| ....|....| ....|....| ....|....|
ETH_00031750               PNVDPTGGPT GAPRRAPCEA PEGPPQGDEG SLPEN-DQIC VAVAIR-MAP GCSIVDNFVL  395
Vbra_16581                 ---------- ---------- ---------- ------KSYC VAIGAS-VGP SCRLVDNIVL  962
Cvel_3471                  ---------- -CLHAGGSVK VSKP------ ---------V LLPSGT-PRE RVRLVDNVVI  1129
SN3_00101300               SRLQPEGEAN HERKVDGCEQ QQEDKEGVEE RLLESPGKYC CTVAVR-GNA DTTLVDNVTL  886
NCLIV_039640               ---------- ---------- ---------- ---------- -----R-ANP ----------  392
TGME49_265870              SSYKRGGHLK GELHANGSSR SSSPADTADM ILLDS-GRYC LTVAVR-GNP LTTVVDSVFL  551
HHA_265870                 SSYKRGGHLK GELHANGSSH SSSPADTADM ILLDS-GRYC LTVAVR-GNP LTTVVDSVVL  551
XP_002768075.P. marinus    ---------- ---------- ---------- ------EPVL VAIAVKYGGA NLRLIDNMWM  325
XP_002767048.P. marinus    ---------- ---------- ---------- ------EPVL VAIAVKYGGA NLRLIDNMWM  325
sp|P31663|PANC_ECOLI       ---------- ---------- ---------- ------KRAV ILVAAW-LGD -ARLIDNKMV  280
sp|P9WIL5|PANC_MYCTU       ---------- ---------- ---------- ---------- LLVAAR-LGT -TRLLDNIAI  285

                                   1390       1400       1410       1420       1430       1440
                           ....|....| ....|....| ....|....| ....|....| ....|....| ....|....|
ETH_00031750               SDTHKFGDLL PPVPNSAEAV PSAAAALLPL HAAPCVLLLR SLSLQLLQKS PPAAAAAAAA  455
Vbra_16581                 EAPKG-DSAQ QRQQQSGWMA SWLGAGK--- ---------- ---------- ----------  988
Cvel_3471                  RVP------- ---------- ---------- ---------- ---------- ----------  1132
SN3_00101300               TDQPQCGDLL PPPLNVPWGR NSAAAKHSFF ---GSFLVLA SLGMHLRYLN EDDVDLLLQA  943
NCLIV_039640               ---------- ---------- ---------- ---------- ---------- ----------  392
TGME49_265870              IPNNR-EDLL PPLFNTPWDH HCVEPKYACL ---PSFLVFS SFGFHLRPLR PSDLKLVAAA  607
HHA_265870                 IPNNR-EDLL PPLFNTPWDH HCVEPKYACL ---PSFLLFS SFGFHLRPLR PSDLKLVVAA  607
XP_002768075.P. marinus    DNQKHA---- ---------- ---------- ---------- ---------- ----------  331
XP_002767048.P. marinus    DNQKHA---- ---------- ---------- ---------- ---------- ----------  331
sp|P31663|PANC_ECOLI       ELA------- ---------- ---------- ---------- ---------- ----------  283
sp|P9WIL5|PANC_MYCTU       EIGTFAGTDR PDGYRAILES HWRN------ ---------- ---------- ----------  309

                                   1450       1460       1470       1480       1490       1500
                           ....|....| ....|....| ....|....| ....|....| ....|....| ....|....|
ETH_00031750               AEWAQTVEE- ---------- ---------- ---------- ---------- ----------  464
Vbra_16581                 ---------- ---------- ---------- ---------- ---------- ----------  988
Cvel_3471                  ---------- ---------- ---------- ---------- ---------- ----------  1132
SN3_00101300               EEEDMEPRQN GPRLLLPPLS PAVGATAACQ RQTEGEKNLG CSTRQQPVAA AGPGTETREP  1003
NCLIV_039640               ---------- ---------- ---------- ---------- ---------- ----------  392
TGME49_265870              EQEDARVES- ---------- ---------- ---------- ---------- ----------  616
HHA_265870                 EQEDARVES- ---------- ---------- ---------- ---------- ----------  616
XP_002768075.P. marinus    ---------- ---------- ---------- ---------- ---------- ----------  331
XP_002767048.P. marinus    ---------- ---------- ---------- ---------- ---------- ----------  331
sp|P31663|PANC_ECOLI       ---------- ---------- ---------- ---------- ---------- ----------  283
sp|P9WIL5|PANC_MYCTU       ---------- ---------- ---------- ---------- ---------- ----------  309

                                   1510       1520       1530       1540       1550       1560
                           ....|....| ....|....| ....|....| ....|....| ....|....| ....|....|
ETH_00031750               ---------- ---------- ---------- ---------- --------LE ALELSDPVVS  476
Vbra_16581                 ---------- ---------- ---------- ---------- ---------- ----------  988
Cvel_3471                  ---------- ---------- ---------- ---------- ---------- ----------  1132
SN3_00101300               LLQQQQRIGV CEGTRARTGE RMHILRWLQQ QRRTLFEVEA AGTRKTLWMK KQGDTAAAGA  1063
NCLIV_039640               ---------- ---------- ---------- ---------- ---------- ----------  392
TGME49_265870              ---------- ---------- ---------- ---------- ---------Q QSAKSAGVAA  627
HHA_265870                 ---------- ---------- ---------- ---------- ---------Q QSAKSTGADA  627
XP_002768075.P. marinus    ---------- ---------- ---------- ---------- ---------- ----------  331
XP_002767048.P. marinus    ---------- ---------- ---------- ---------- ---------- ----------  331
sp|P31663|PANC_ECOLI       ---------- ---------- ---------- ---------- ---------- ----------  283
sp|P9WIL5|PANC_MYCTU       ---------- ---------- ---------- ---------- ---------- ----------  309

                                   1570       1580       1590       1600       1610       1620
                           ....|....| ....|....| ....|....| ....|....| ....|....| ....|....|
ETH_00031750               SLSSAERQKA WPWASAAAAA AAAP------ ---------- ---------- ----------  500
Vbra_16581                 ---------- ---------- ---------- ---------- ---------- ----------  988
Cvel_3471                  ---------- ---------- ---------- ---------- ---------- ----------  1132
SN3_00101300               AEQSCVRPCA QQSSQVHSPA TSGSGSTFQN SSNVCRVRAV GLFKYDLMDE CSSSSNETAR  1123
NCLIV_039640               ---------- ---------- ---------- ---------- ---------- ----------  392
TGME49_265870              PETCCTRTFT WMHAGALSPT STHD------ ------VWAL GLFKRRL--- ----------  662
HHA_265870                 PETCCTRTFT WMHAGALSPT STHD------ ------VWAL GLFKRRL--- ----------  662
XP_002768075.P. marinus    ---------- ---------- ---------- ---------- ---------- ----------  331
XP_002767048.P. marinus    ---------- ---------- ---------- ---------- ---------- ----------  331
sp|P31663|PANC_ECOLI       ---------- ---------- ---------- ---------- ---------- ----------  283
sp|P9WIL5|PANC_MYCTU       ---------- ---------- ---------- ---------- ---------- ----------  309

                                   1630       1640       1650       1660       1670       1680
                           ....|....| ....|....| ....|....| ....|....| ....|....| ....|....|
ETH_00031750               ---------- ---------- ---------- ---------- ---------- ----------  500
Vbra_16581                 ---------- ---------- ---------- ---------- ---------- ----------  988
Cvel_3471                  ---------- ---------- ---------- ---------- ---------- ----------  1132
SN3_00101300               LTSHCSDRAA LEADRISVAS RGGADASIQS KDTRRREGRH VAAVGDTSAN RTALHSGGRT  1183
NCLIV_039640               ---------- ---------- ---------- ---------- ---------- ----------  392
TGME49_265870              ---------- ---------- ---------- ---------- -----DTARN TVDFSSDTQV  677
HHA_265870                 ---------- ---------- ---------- ---------- -----DSARN TVDVSSDTQV  677
XP_002768075.P. marinus    ---------- ---------- ---------- ---------- ---------- ----------  331
XP_002767048.P. marinus    ---------- ---------- ---------- ---------- ---------- ----------  331
sp|P31663|PANC_ECOLI       ---------- ---------- ---------- ---------- ---------- ----------  283
sp|P9WIL5|PANC_MYCTU       ---------- ---------- ---------- ---------- ---------- ----------  309

                                   1690       1700       1710       1720       1730       1740
                           ....|....| ....|....| ....|....| ....|....| ....|....| ....|....|
ETH_00031750               ---------- ---------- ---------- ---------- --SSYFLLRR LRCTDSPGEL  518
Vbra_16581                 ---------- ---------- ---------- ---------- ---------- ----------  988
Cvel_3471                  ---------- ---------- ---------- ---------- ---------- ----------  1132
SN3_00101300               PAERDTCLSG AADRCSDSGN PRETTAKSGR FKRRERLIGY VWAEEYAGEE QGGPPQKRER  1243
NCLIV_039640               ---------- ---------- ---------- ---------- ---------- ----------  392
TGME49_265870              PDGNEV---- ---------- ---------- ------LIGY IWAD------ --------AE  693
HHA_265870                 PDGNEV---- ---------- ---------- ------LIGY IWAD------ --------AE  693
XP_002768075.P. marinus    ---------- ---------- ---------- ---------- ---------- ----------  331
XP_002767048.P. marinus    ---------- ---------- ---------- ---------- ---------- ----------  331
sp|P31663|PANC_ECOLI       ---------- ---------- ---------- ---------- ---------- ----------  283
sp|P9WIL5|PANC_MYCTU       ---------- ---------- ---------- ---------- ---------- ----------  309

                                   1750       1760       1770       1780       1790       1800
                           ....|....| ....|....| ....|....| ....|....| ....|....| ....|....|
ETH_00031750               VAYLAFTRVQ NKSICIQRIF VAPKERRKGI AEAFFCTCML LLQQQLLQQQ LQQQQASNEQ  578
Vbra_16581                 ---------- ---------- ---------- ---------- ---------- ----------  988
Cvel_3471                  ---------- ---------- ---------- ---------- ---------- ----------  1132
SN3_00101300               KEGCSASGNV CRVLRVNRVF LKRERRRQTL GTHLLRAFIA ATYKQLLEEQ QQAVSVAPRH  1303
NCLIV_039640               ---------- ---------- ---------- ---------- ---------- ----------  392
TGME49_265870              TPRAAAAAGG ASPWRLRRGF ITRARRRRTF GTHLLKAFLA TLYKEKVSSS RTA-------  746
HHA_265870                 TPHAAAAAGG ASPWRLRRGF ITRARRRRTF GTHLLKAFLA TLYREKVSSN RTA-------  746
XP_002768075.P. marinus    ---------- ---------- ---------- ---------- ---------- ----------  331
XP_002767048.P. marinus    ---------- ---------- ---------- ---------- ---------- ----------  331
sp|P31663|PANC_ECOLI       ---------- ---------- ---------- ---------- ---------- ----------  283
sp|P9WIL5|PANC_MYCTU       ---------- ---------- ---------- ---------- ---------- ----------  309

                                   1810       1820       1830       1840       1850       1860
                           ....|....| ....|....| ....|....| ....|....| ....|....| ....|....|
ETH_00031750               ---------- ---------- ---------- ---------- ---------- ----------  578
Vbra_16581                 ---------- ---------- ---------- ---------- ---------- ----------  988
Cvel_3471                  ---------- ---------- ---------- ---------- ---------- ----------  1132
SN3_00101300               HRPPSAPVLH QRQQQEAVLK TAGHTGPTVK GRPQECEDRE AVVCKIRRTE EEQEMQEDAI  1363
NCLIV_039640               ---------- ---------- ---------- ---------- ---------- ----------  392
TGME49_265870              ---------- ---------- ---------- ---------- ---------- ----------  746
HHA_265870                 ---------- ---------- ---------- ---------- ---------- ----------  746
XP_002768075.P. marinus    ---------- ---------- ---------- ---------- ---------- ----------  331
XP_002767048.P. marinus    ---------- ---------- ---------- ---------- ---------- ----------  331
sp|P31663|PANC_ECOLI       ---------- ---------- ---------- ---------- ---------- ----------  283
sp|P9WIL5|PANC_MYCTU       ---------- ---------- ---------- ---------- ---------- ----------  309

                                   1870       1880       1890       1900       1910       1920
                           ....|....| ....|....| ....|....| ....|....| ....|....| ....|....|
ETH_00031750               ---------- ---------- ---------- ---------- ---------- ---QVTCVVP  585
Vbra_16581                 ---------- ---------- ---------- ---------- ---------- ----------  988
Cvel_3471                  ---------- ---------- ---------- ---------- ---------- ----------  1132
SN3_00101300               TEATETRKSR ERGKLETLCV EGETESTLEG GATEGKEWKK SKHEMEGEIS CTVQLENVPH  1423
NCLIV_039640               ---------- ---------- ---------- ---------- ---------- ----------  392
TGME49_265870              --------SR EV-------- ---------- ---------- ---------- ---QGDPMPA  757
HHA_265870                 --------SR EV-------- ---------- ---------- ---------- ---QGDPMPA  757
XP_002768075.P. marinus    ---------- ---------- ---------- ---------- ---------- ----------  331
XP_002767048.P. marinus    ---------- ---------- ---------- ---------- ---------- ----------  331
sp|P31663|PANC_ECOLI       ---------- ---------- ---------- ---------- ---------- ----------  283
sp|P9WIL5|PANC_MYCTU       ---------- ---------- ---------- ---------- ---------- ----------  309

                                   1930       1940       1950       1960       1970
                           ....|....| ....|....| ....|....| ....|....| ....|....| ...
ETH_00031750               QLALKVMEKS FGFTPEGGPH EGGPHQGGPH EGGTPTVAVS LELNALAQRL LRP 638
Vbra_16581                 ---------- ---------- ---------- ---------- ---------- --- 988
Cvel_3471                  ---------- ---------- ---------- ---------- ---------- --- 1132
SN3_00101300               YLVGVLREGG FCVDTAAAET TRAQTYKGSC CCMHVSLAQW YNAFVLNANV --- 1473
NCLIV_039640               ---------- ---------- ---------- ---------- ---------- --- 392
TGME49_265870              WLAPAVSECG FSL------- LDSTNLSSGE CRLHLNLDKW FRTYVTDLRL --- 800
HHA_265870                 WLAPAVSECG FSL------- MESTNLSSGE CRLHLNLDKW FRANVTDLGL --- 800
XP_002768075.P. marinus    ---------- ---------- ---------- ---------- ---------- --- 331
XP_002767048.P. marinus    ---------- ---------- ---------- ---------- ---------- --- 331
sp|P31663|PANC_ECOLI       ---------- ---------- ---------- ---------- ---------- --- 283
sp|P9WIL5|PANC_MYCTU       ---------- ---------- ---------- ---------- ---------- --- 309
```
