## Supplementary File 2 for "Pantothenate biosynthesis is critical for chronic infection by the neurotropic parasite *Toxoplasma gondii*"

**Supplementary Material and Methods – liquid chromatography conditions and mass spectrometry parameters**

**Reversed-phase UPLC-MS conditions**

Quantification of Pan, PPan-Cys, deP-CoA and CoA (method A) was performed on the Q Exactive Plus analytical platform as follows: 10 μl of sample (6°C) were injected and eluted with a tailored gradient: 0%B (1 min), 5-25%B in 7 min, 25-40%B in 1 min and hold 3 min, wash at 95%B (total runtime of 20 min). Here and in all other methods, blanks (ultrapure water or 80% acetonitrile for HILIC analyses) were injected after every sample to avoid carry-over. Eluent A was 50 mM ammonium acetate buffer (pH 6.9) and eluent B was acetonitrile. Flow rate and column oven temperature were respectively of 250 µl/min and 30 °C. The mass spectrometer was operated in positive polarity with a heated electrospray ionization (HESI-II) probe. Electrospray voltage was set to 3500 V; the sheath gas, auxiliary gas and sweep gas were set, respectively, to 46, 11 and 2 (arbitrary units, nitrogen). The auxiliary gas heater and transfer capillary temperatures were of 350 °C and 300 °C. PRM experiments were looped for each precursor with an isolation window of 0.7u, a normalized collision energy of 30 eV. All experiments were acquired in profile mode at 17.5 k resolution with an AGC of 1e6 and a fill time of 50 ms (1 µscan).

For labelling experiments (method B - Fig. 5f), 5 μl of sample (6°C) were injected and separated with a linear gradient from 0%B (1 min hold) to 95%B in 14 min (total runtime of 20 min after column wash and reconditioning). Eluent A was 5 mM ammonium formate with 0.25% formic acid (pH 2.8) and eluent B was acetonitrile. Flow rate and column oven temperature were respectively of 250 µl/min and 35 °C. Mass spectrometry experiments were a full scan MS looped with PRM (FS-PRM) experiments for each precursor (isol. window of 0.7u - NCE = 25). Electrospray interface settings were as described above, and MS data was acquired in profile mode at 35 k resolution with an AGC of 1e6 and a fill time of 100 ms (1 µscan).

For the QTRAP 3200 analytical platform, 10 μl of sample (8°C) were injected and eluted with the same gradient as for method A with the Q Exactive Plus instrumentation. The mass spectrometer was operated in positive polarity with a TurboV ion source fitted with a TurboIonSpray probe (SCIEX). The electrospray voltage was set to 5500 V, the nebulizer gas and auxiliary gas (nitrogen) were set to 40 and 50 psi, respectively. The auxiliary gas heater was set to 500 °C and the curtain plate (CUR), entrance lens (EP) and collision cell exit (CXP) potentials were of 10 V. The declustering potential (DP) was set to 70 V. The MRM transitions of the precursor ion (Q1) and product ion (Q3), as well as the collision energy (CE) are summarized in the table (Q1 and Q3 were operated at unit resolution):

| **Name** | **Q1 (*m/z*)** | **Q3 (*m/z*)** | **Dwell (ms)** | **CE (eV)** |
| --- | --- | --- | --- | --- |
| β-Ala_1 | 90.1 | 44.0 | 30 | 40 |
| ^13^C_3_/^15^N-β-Ala_1 | 94.1 | 47.0 | 30 | 40 |
| β-Ala_2 | 90.1 | 55.0 | 30 | 30 |
| ^13^C_3_/^15^N-β-Ala_2 | 94.1 | 58.0 | 30 | 30 |
| Panthotenate_1 | 220.1 | 90.0 | 30 | 35 |
| ^13^C_3_/^15^N-panthotenate_1 | 224.1 | 94.2 | 30 | 35 |
| Panthotenate_2 | 220.1 | 71.9 | 30 | 45 |
| ^13^C_3_/^15^N-panthotenate_2 | 224.1 | 75.9 | 30 | 45 |
| CoA_1 | 768.1 | 261.0 | 30 | 26 |
| ^13^C_3_/^15^N-CoA_1 | 772.1 | 265.0 | 30 | 26 |
| CoA_2 | 768.1 | 428.0 | 30 | 23 |
| ^13^C_3_/^15^N-CoA_2 | 772.1 | 428.0 | 30 | 23 |
| Acetyl-CoA_1 | 810.1 | 303.0 | 30 | 27 |
| ^13^C_3_/^15^N-acetyl-CoA_1 | 814.1 | 307.0 | 30 | 27 |
| Acetyl-CoA_2 | 810.1 | 428.0 | 30 | 24 |
| ^13^C_3_/^15^N-acetyl-CoA_2 | 814.1 | 428.0 | 30 | 24 |

For the analyses performed with the QTRAP 6500, the following UPLC-MS conditions were used: 8 μl of sample (6°C) were injected and eluted with the same gradient and column as described for method A. The mass spectrometer was operated with the same settings as previously mentioned except the curtain plate (CUR) and collision cell exit (CXP) potentials set to 30 V and 8 V respectively. The declustering potential (DP) was of 90 V. The set of MRM transitions was the same with the addition of two transitions to monitor dephospho-CoA:

| **Name** | **Q1 (*m/z*)** | **Q3 (*m/z*)** | **Dwell (ms)** | **CE (eV)** |
| --- | --- | --- | --- | --- |
| Dephospho-CoA_1 | 688.2 | 261.1 | 30 | 26 |
| Dephospho-CoA_2 | 688.2 | 348.1 | 30 | 23 |

**HILIC UPLC-MS conditions**

HILIC analyses were carried out on the Q Exactive Plus analytical platform as follows: 5 μl of sample (6°C) were injected and eluted with a gradient adapted ^71^: 100%B held for 6 min and stepped down to 94.1%B, 94.1-82.4%B in 4 min, 82.4-70.6%B in 2 min and back to 100%B (total runtime of 20 min). Eluent A was 10 mM ammonium formate with 0.15% formic acid in water and eluent B was 10 mM ammonium formate with 0.15% formic acid in acetonitrile-water (85:15, v/v). Flow rate and column oven temperature were respectively of 400 µl/min and 40 °C. The mass spectrometer was operated in positive or negative polarity with a heated electrospray ionization (HESI-II) probe. Electrospray settings were as aforementioned. A full MS experiment was followed with PRM experiments looped for each precursor with an isolation window of 0.7u, a normalized collision energy of 30 eV. All experiments were acquired in profile mode at 17.5 k resolution with an AGC of 1e6 and a fill time of 50 ms (1 µscan).
